# The functional impact of LGI1 autoantibodies on human CA3 pyramidal neurons

**DOI:** 10.1101/2024.10.28.620296

**Authors:** Laura Monni, Hans-Christian Kornau, Alice Podestà, Alexander Stumpf, Thilo Kalbhenn, Matthias Simon, Thomas Sauvigny, Julia Onken, Harald Prüss, Henrik Alle, Jörg R.P. Geiger, Martin Holtkamp, Dietmar Schmitz, Pawel Fidzinski

## Abstract

Autoantibodies against leucine-rich glioma inactivated 1 protein (LGI1 mAb) lead to limbic encephalitis characterized by seizures and memory deficits. While animal models provide insights into mechanisms of LGI1 mAb action, species-specific confirmation is lacking. In this study, we investigated the effects of patient-derived LGI1 mAb on human CA3 neurons using cultured *ex vivo* slices. Analysis of intrinsic properties and morphology indicated functional integrity of these neurons under incubation conditions. Human CA3 neurons received spontaneous excitatory currents with large amplitudes and frequencies, suggestive of “giant” AMPA currents. In slices exposed to LGI1 mAb, human CA3 neurons displayed increased neuronal spike frequency, mirroring effects observed with the K_v_1.1 channel blocker DTX-K. This increase likely resulted from decreased K_v_1.1 channel activity at the axonal initial segment, as indicated by alterations in action potential properties. A detailed analysis revealed differences between LGI1 mAb and DTX-K effects on action potential properties, suggesting distinct mechanisms of action and emphasizing the need for further exploration of downstream pathways. Our findings underscore the importance of species-specific confirmatory studies of disease mechanisms and highlight the potential of human hippocampal slice cultures as a translational model for investigation of disease mechanisms beyond epilepsy, including the effects of pharmacological compounds and autoantibodies.

**Significance:** This study advances our understanding of how autoantibodies against the LGI1 protein, known to cause limbic encephalitis, impact human neurons. By using cultured slices of human hippocampus derived from epilepsy’s surgical resections, we were able to observe the direct effects of these autoantibodies on neurons, specifically CA3 pyramidal cells. Our findings reveal that the autoantibodies increase neuronal activity, similar to what is seen with potassium channel blockers and in animal models. This work emphasizes the importance of studying living tissue from the human brain to confirm disease mechanisms, and demonstrates the potential of using human brain slices as a model for exploring not only epilepsy but also other neurological diseases and drug effects.

## Introduction

Autoimmune encephalitis (AE) refers to a diverse group of neurological disorders characterized by the production of autoantibodies against neuronal antigens. Among these, autoantibodies targeting leucine-rich glioma-inactivated 1 (LGI1) protein are particularly significant due to their impact on the limbic system^1,2^, leading to epilepsy and cognitive deficits^2–5^. The disease manifests variably, ranging from mild faciobrachial dystonic seizures to severe, treatment-refractory seizures. While immunotherapy often alleviates symptoms, patients frequently suffer from enduring cognitive impairments and seizures, resulting in long-term disability^6,7^.

LGI1 is a component of a multiprotein complex at excitatory synapses and the axon initial segment (AIS), facilitating transsynaptic interactions and modulating neuronal intrinsic excitability^8–10^. In addition to leucine-rich repeat (LRR) and epitempin (EPTP) domains of the LGI1 protein, the complex contains ADAM22/23 receptors and the PSD-95 family associated guanylate kinases (MAGUKs)^8,11^. These components together regulate the expression and activity of postsynaptic α-amino-3-hydroxy-5-methyl-4-isoxazolepropionic acid (AMPA) receptors, which mediate rapid excitatory neurotransmission and contribute to single-cell and population activity in the brain^12,13^. Additionally, the components of the LGI1 protein complex modulate presynaptic K_v_1.1 voltage-gated potassium channel expression and activity, in particular at the AIS^9,11^. K_v_1.1 channels mediate the D-type current, a key determinant of neuronal excitability^14,15^ and intrinsic homeostatic plasticity in the hippocampal CA3 microcircuit^9,10,16^. The CA3 network is essential for hippocampus-mediated cognitive functions such as learning, memory consolidation, and spatial navigation^17^.

Recent technological advancements have enabled recombinant cloning and production of patient-derived autoantibodies in large quantities, facilitating their functional characterization in animal models^18,19^. Specifically, LGI1 autoantibodies have been investigated in rodent models to elucidate their mechanisms of action^20–25^. Functionally, LGI1 autoantibodies increase intrinsic excitability and presynaptic release probability in murine CA3 neurons and mossy fibers^21,23,26,27^. Current evidence indicates that these effects are mediated by the impairment of K_v_1.1 channel function. While animal models provide valuable insights into pathomechanisms, they have significant limitations. Recent studies have revealed differences between rodents and humans in neuronal cell types and architecture, dendritic properties, and synaptic plasticity^28–36^. These discrepancies contribute to the translational gap between bench research and clinical application^37–42^, emphasizing the need for more suitable models to investigate human disease. Importantly, specific cellular mechanisms of LGI1 autoantibodies in the human hippocampus remain largely unexplored.

Recognizing the limitations associated with rodent models, particularly species-specific differences^43–45^, we aimed to investigate the effects of LGI1 autoantibodies on CA3 pyramidal cells and the CA3 network in organotypic hippocampal slice cultures from temporal lobe epilepsy patients undergoing hippocampal resection. Specifically, we characterized the biophysical properties of human CA3 pyramidal neurons *ex vivo* and examined how these properties are altered in slices incubated in the presence of human-derived LGI1 monoclonal antibodies (LGI1 mAb).

## Materials and Methods

### Human tissue transport and slice preparation

Human hippocampal tissue from epilepsy surgical resection was collected from 14 patients diagnosed with pharmacoresistant temporal lobe epilepsy (11 males, 3 females; 13-60 years old; see supplementary table 1 for details). All patients provided written consent. The study adhered to the Declaration of Helsinki and was approved by the Ethics Committees of Charité University Medicine Berlin, Westfalen-Lippe for Bielefeld University, and University Medical Center Hamburg-Eppendorf (EA2/111/14, EA2/064/22 and 2020/517/f/S, respectively).

**Table 1:** analyzed passive properties of human CA3 pyramidal neurons after 0-3 h from recovery and 18-24 h incubation in untreated control conditions. Data are expressed as mean ± SD.

| Passive properties |  | 0-3 h | 18-24 h |
| --- | --- | --- | --- |
| RMP (mV) | Mean | -54.3 | -60.4 |
|  | SD | 4.9 | 5.4 |
| Input R (MΩ) | Mean | 134.1 | 105.6 |
|  | SD | 27.5 | 43.5 |
| Rheobase (pA) | Mean | 134.7 | 309.1 |
|  | SD | 88.6 | 103.3 |
| Membrane Tau (ms) | Mean | 57.1 | 39.9 |
|  | SD | 23.7 | 15.0 |
| Voltage Sag (mV) | Mean | -3.3 | -3.7 |
|  | SD | 0.5 | 1.9 |
| Sag ratio | Mean | 0.12 | 0.18 |
|  | SD | 0.02 | 0.08 |

Resected hippocampal tissue was sourced from three locations: local neurosurgical department at the Charité University Medicine Berlin, Evangelisches Klinikum Bethel of the Bielefeld University, and University Medical Center Hamburg-Eppendorf. Immediately after resection, part of the anterior hippocampus was transferred *en bloc* in a carbogenated (95% O_2_, 5% CO_2_), ice-cold sterile-filtered choline-based solution (Figure 1A-B). The sample was transported to our laboratory in Berlin either directly or via train in gas-tight sterile bottles secured in a styrofoam box with ice. Transport time from the operating theater to the laboratory ranged from 15 min to 5 hours without noticeable differences in tissue quality. As resected hippocampi from epileptic patients frequently show morphological alterations and varying degrees of sclerosis, in particular in the CA1 region^46,47^, we visually inspected all specimens and included only tissue that did not show evident signs of neuronal loss and sclerosis in the CA3 subregion. Evident signs included the marked reduction in the number of pyramidal neurons, thinning or loss of neurons in the dentate gyrus granule cell layer, as well as visible shrinkage of the hippocampal structure^46^.

**Figure 1:**
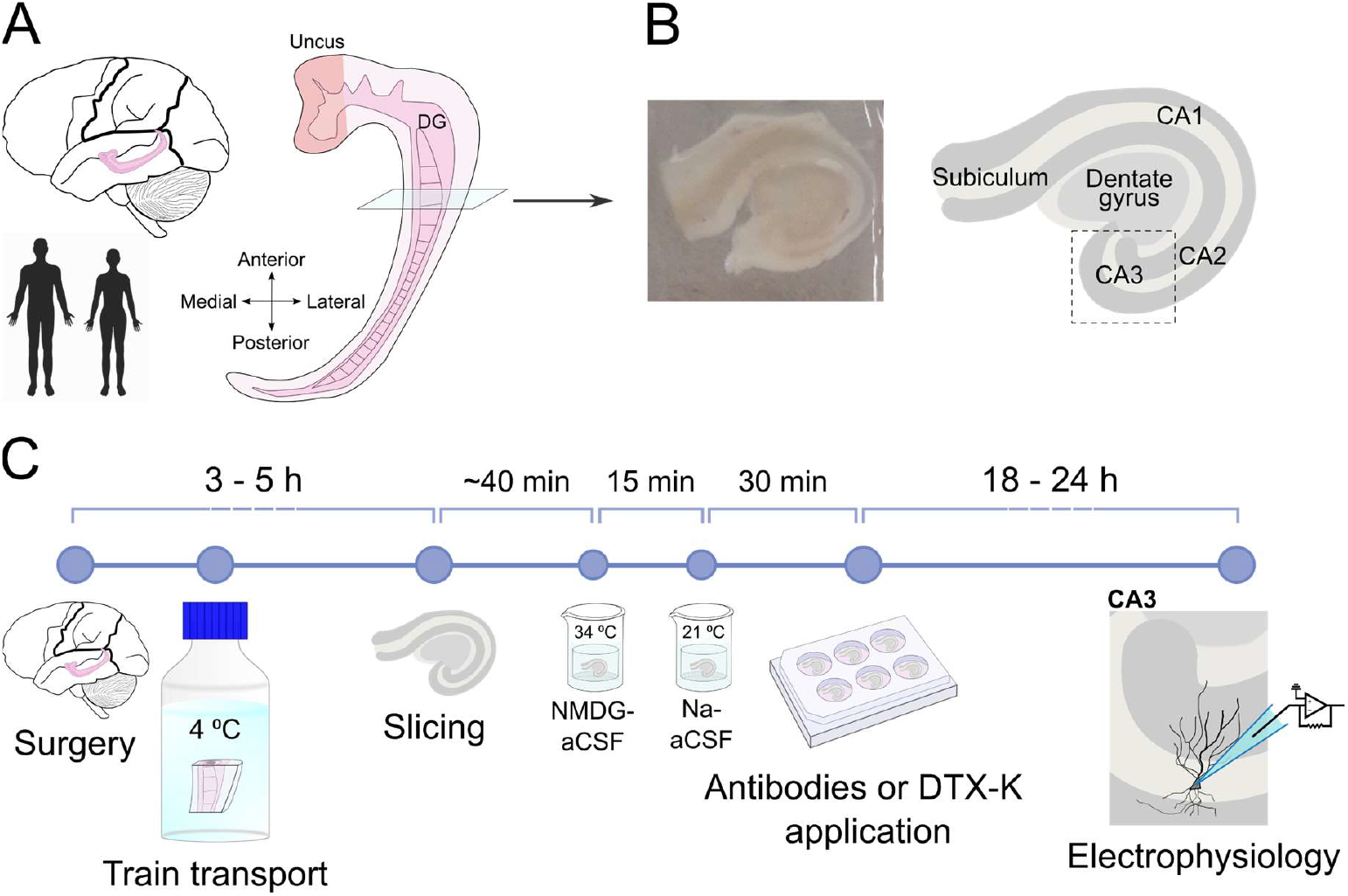
Experimental setting. **A)** Schematic view of the human hippocampus. All samples were taken from comparable mid-anterior hippocampal body from female and male epileptic patients. **B)** Left, representative image of a typical 300 μm-thick acute slice obtained via vibratome-cutting. Right, schematic of the same slice with main hippocampal subregions: the subiculum, the cornus ammonis (CA) 1, CA2, CA3, and the dentate gyrus. Dashed line indicates location of CA3 region. **C)** Timeline showing the experimental procedure (for details, see Methods).

As previously reported^48^, hippocampal slices were cut into 300 μm-thick slices using a vibratome (Leica VT1200S, Wetzlar, Germany) in ice-cold and carbogenated slicing solution. Only slices containing all hippocampal subregions (subiculum, cornus ammonis (CA) 1, CA2, CA3 and dentate gyrus (DG)) were used (Figure 1B). Artificial cerebrospinal fluid (aCSF) containing choline as substitution of sodium ions was used both for hippocampal tissue transport and for slice-cutting. Choline-aCSF (305 ± 5 mOsm/l, pH 7.4) contained (in mM): choline chloride (110), Na-L-ascorbate (11.6), MgCl_2_ (7), Na-pyruvate (3.1), KCl (2.5), NaH_2_PO_4_ (1.25), NaHCO_3_ (26), CaCl_2_ (0.5), glucose (10)^32^.

### Human hippocampal slice culture

Hippocampal slices were processed as previously reported^48–50^ with modifications. Immediately after slicing, slices were placed in a submerged chamber filled with N-methyl d-glucamine (NMDG)-aCSF at 34 °C for 15 min for recovery. NMDG-aCSF contained (in mM): HEPES (20), Na-pyruvate (3), taurine (0.01), thiourea (2), glucose (25), myo-inositol (3), NaHCO_3_ (30), CaCl_2_ (0.5), MgSO_4_ (10), KCl (2.5), NaH_2_PO_4_ (1.25), HCl (92), NMDG (92), L-ascorbic acid (5), N-acetyl-L-cysteine (12). Upon recovery, slices were then transferred to a submerged chamber containing Na-aCSF at room temperature for 30 min. Na-aCSF contained (in mM): HEPES (20), Na-pyruvate (3), taurine (0.01), thiourea (2), glucose (25), myo-inositol (3), NaHCO_3_ (30), CaCl_2_ (2), MgSO_4_ (2), KCl (2.5), NaH_2_PO_4_ (1.25), NaCl (92), L-ascorbic acid (5), N-acetyl-L-cysteine (12). Both recovery solutions were controlled for pH (adjusted to 7.3-7.4) and osmolarity (300 ± 5 mOsm/l). All solutions were prepared and sterile-filtered on the same day or the day before the surgical hippocampal resection. All procedures were conducted under near-sterile conditions.

After recovery, slices were cultured in individual 0.4 μm-inserts placed in a 6-well plates (Millipore, Bedford, MA, United States). Each well contained 1 ml of culture medium containing (in mM): MEM eagle powder (8.4 g/L), L-ascorbic acid (1), CaCl_2_ (1), MgSO_4_ (2), HEPES (30), NaHCO_3_ (15), glucose (13), tris base (1), glutamine (0.5). The medium was additionally enriched with: 0.2% penicillin/streptavidin antibiotics, 1 mg/L bovine pancreas insulin and 20% heat-inactivated horse serum^50^. Osmolarity and pH of culture medium were adjusted to 300 ± 5 mOsm/l and 7.3-7.4, respectively. The protocol and timeline are illustrated in figure 1C.

### Application of LGI1 monoclonal antibodies and dendrotoxin-K

Human monoclonal antibodies AB060-110 (LGI1 mAb) and AB060-154 (Ab control; an antibody not binding to LGI1 protein) derived from single CSF cells of a limbic encephalitis patient^21^, were expressed in HEK293T cells, purified from cell culture supernatants using Protein G Sepharose 4 Fast Flow (GE Healthcare) and dialyzed against PBS buffer. Their concentrations were determined using a Human IgG ELISA development kit (Mabtech, Sweden).

The functional effects of monoclonal LGI1 antibodies were assessed in individual cultured slices after 18-24 h of antibody exposure. On the first day in culture (DIV0), slices were incubated with LGI1 mAb or Ab control at a final concentration of 6 μg/ml in the culture medium. Dendrotoxin-K (DTX-K) at 100 nM, a selective blocker of K_v_1.1 channels, was used as pharmacological control. An untreated slice served as a control for each replicate.

### Electrophysiological recordings

Patch-clamp recordings in whole-cell configuration and local field potential (LFP) recordings were performed in human hippocampal slices after 18-24 h in culture. All CA3 pyramidal neurons were visually identified. In a subset of slices, patch-clamp recordings were conducted 0-3h after slicing and recovery in order to assess the intrinsic excitability under acute conditions. Borosilicate pipettes (Science Products, Hofheim, Germany; 1.5 mm outer diameter) were pulled with a vertical puller (PC-10, Narishige, Tokyo, Japan; electrode resistance 3-6 MΩ for patch-clamp, or 1-2 MΩ for LFP recordings) and filled with internal solution or 154 mM NaCl for extracellular recordings. Signals were amplified by a Multiclamp 700B Amplifier, sampled at 20 kHz, low-pass filtered at 10 kHz (current clamp) or 2 kHz (LFP recordings and voltage clamp), digitized by a Digidata1550 interface and processed by PClamp10 software (RRID: SCR_011323) (all by Molecular Devices, Sunnyvale, CA, USA). The quality criteria for inclusion of patch-clamp recordings were a > 1 GΩ seal prior to break-in stable series resistance of < 20 MΩ with < 20% change at the end of the recording, and a membrane/series resistance ratio > 8.

Prior to recordings, slices were individually transferred to a modified submerged-type recording chamber characterized by a high flow rate (10 ml/min)^51^ and perfused with carbogenated aCSF (32 °C, 305 ± 5 mOsm/l, pH 7.4) containing (in mM): NaCl (129), NaH_2_PO_4_ (1.25), CaCl_2_ (1.6), KCl (3.0), MgSO_4_ (1.8), NaHCO_3_ (21), glucose (10). Intrinsic excitability of CA3 pyramidal neurons was measured by input-output curves in current clamp configuration, manually holding the membrane potential at -70 mV by continuous current injection. For this set of experiments, we used K-gluconate-based internal solution containing (in mM): K-gluconate (130), HEPES (10), EGTA (0.3), MgCl_2_ (2), MgATP (4), NaGTP (0.3), osmolarity 290 ± 5 mOsm, pH 7.2-7.4. Liquid junction potential was -14 mV and was not corrected for. Current clamp recordings of action potential firing were performed in absence of synaptic input obtained by pharmacological blockade of synaptic transmission with NBQX (10 μM) to block α-amino-3-hydroxy-5-methyl-4-isoxazolepropionic acid receptors (AMPA), APV (50 μM) to block N-methyl-D-aspartate receptors (NMDA), and Gabazine (1 μM) to block γ-aminobutyric acid receptors (GABA_A_). Input-output curves were obtained by recording the membrane voltage changes in response to increasing current injections lasting 500 ms, incremented by 50 pA (from -400 pA to 800 pA).

A subset of CA3 pyramidal neurons from untreated control slices was used to record spontaneous excitatory currents recorded and filled with biocytin-containing internal solution for at least 20 minutes for morphological reconstruction. AMPA miniature excitatory postsynaptic currents (mEPSCs) were recorded in voltage clamp configuration at a membrane potential held at -70 mV. Only neurons displaying a stable holding current ≥ -300 pA were included in the study. The internal solution was caesium-based and contained (in mM): CsCl (130), MgCl_2_ (2), HEPES (10), MgATP (2), EGTA (0.2), QX-314 (5), 0.2% biocytin, osmolarity 290 ± 5 mOsm, pH 7.2-7.4. mEPSCs were isolated with the Na^+^ channels blocker tetrodotoxin (TTX, 1 μM). Gabazine (1 μM) and APV (50 μM) were added to the external aCSF to block GABA_A_ and NMDA receptors, respectively.

Spontaneous network activity was measured by LFP recordings in CA3 pyramidal layer for at least 20 minutes. Slices were perfused with aCSF containing (in mM): NaCl (124), NaH_2_PO_4_ (1.25), CaCl_2_ (1.6), KCl (5.0), MgSO_4_ (1.8), NaHCO_3_ (21), glucose (10). Due to technical reasons, *in vitro* experiments were not blinded, however, electrophysiology data processing and statistical analysis were conducted in a blinded fashion.

### Electrophysiology data processing

Electrophysiological data were processed and analyzed using PClamp 10.7 software (Molecular Devices, USA) for raw data processing and analysis of local field potential (LFP) recordings and Easy Electrophysiology software (Easy Electrophysiology Ltd., UK; RRID: SCR_021190) for analyzing neuronal firing activity, action potential properties, and miniature excitatory postsynaptic currents (mEPSCs).

#### Passive properties of CA3 pyramidal neurons

Resting membrane potential (RMP) was assessed within the first minute after establishing whole-cell configuration. Averaged input resistance was calculated using responses to -100, -50, 50 and 100 pA current steps. Rheobase was defined as the minimal current required to trigger at least one action potential (AP). Membrane time constant was estimated by averaging the time constants (T-values) from exponential fits to the first 150 ms of voltage responses to aforementioned current steps. Voltage sag amplitude was measured as the voltage difference between the peak of the hyperpolarized deflection and the steady-state membrane potential in response to a hyperpolarizing -200 pA current step. Sag ratio was calculated by dividing the voltage sag with the hyperpolarized peak voltage. Refer to Figure 2B for a schematic representation of these calculations.

**Figure 2:**
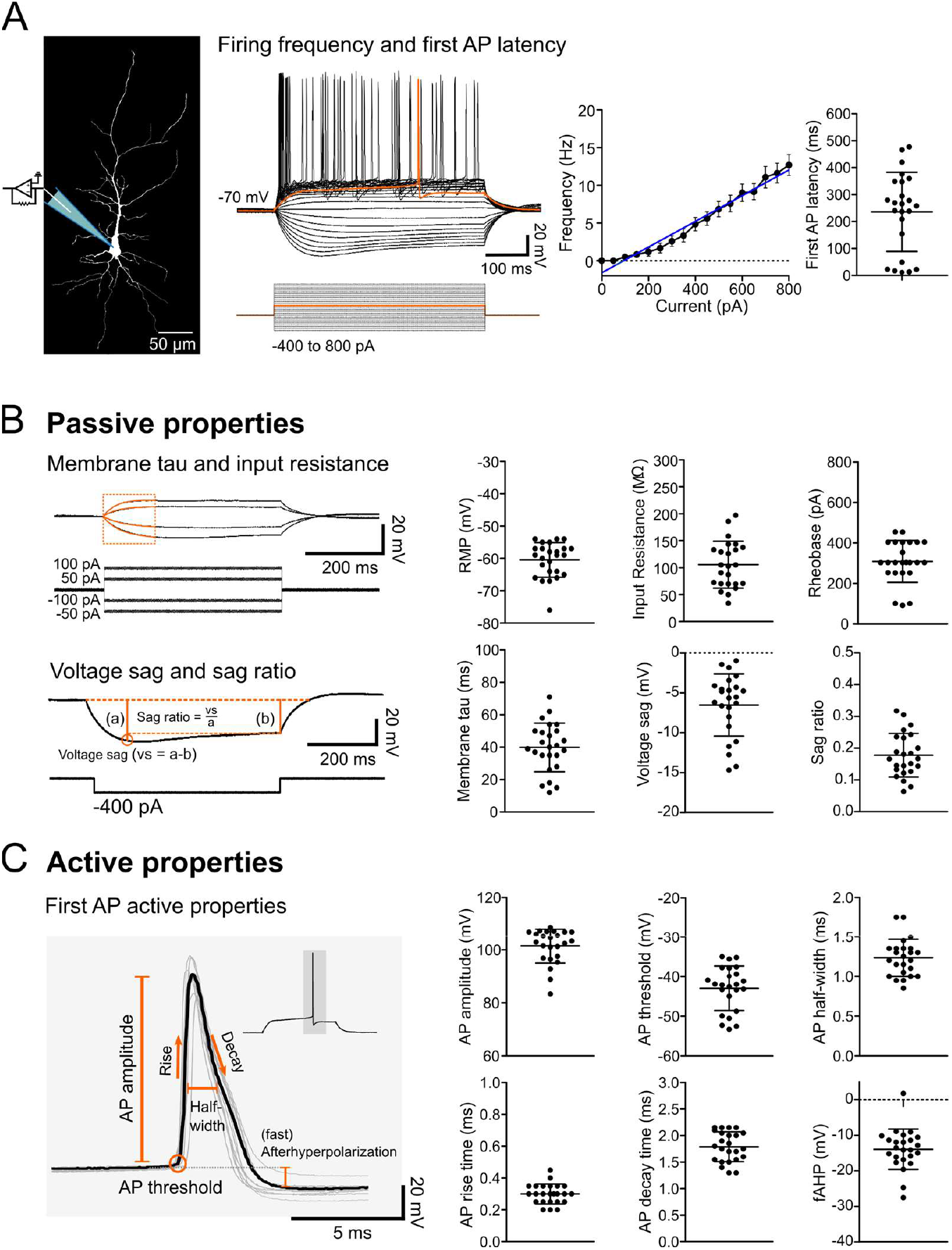
Biophysical properties of human CA3 pyramidal cell properties. **A)** Left: biocytin-filled pyramidal neuron from human hippocampal CA3 region. Middle: representative voltage traces induced by increasing hyperpolarizing and depolarizing current injections (from -400 to 800 pA, 50 pA steps). The first action potential (AP) evoked at the rheobase is highlighted in orange. Right: neuronal firing frequency plotted against current injections showing the input-output relation of CA3 pyramidal cells (shown as mean ± SEM, n = 24 from 9 patients, blue line: r^2^=0.9723) and a scatter plot showing the first AP latency. **B)** Left: experimental steps used to determine passive membrane properties. Right: scatter plots depicting passive properties: resting membrane potential (RMP), input resistance, rheobase, membrane time constant (tau), voltage sag, and sag ratio. **C)** Left: APs recorded at rheobase and employed to quantify active properties. Thick line indicates the average AP of all traces. Right: analysis of active properties: AP amplitude, threshold, half-width, rise time, decay time and fast afterhyperpolarization (fAHP), shown as scatter plots. **B)-C)** Each point indicates an individual cell, and data are shown as mean ± SD (n = 24 from 9 patients).

#### Active properties of action potentials

Neuronal firing properties were assessed by the number of APs evoked by increasing current injections. Six key properties were calculated from the first AP elicited at the rheobase: AP threshold was defined as membrane potential at which the first derivative of voltage exceeded 20 mV/ms; AP amplitude was considered as the peak voltage of the AP from the AP threshold; AP half-width was calculated as time between the two half-amplitude points of AP rise and decay; AP rise (and decay) times were calculated as time (in ms) between 10 and 90% (or 90 and 10%) of the AP amplitude, respectively; fast afterhyperpolarization was calculated as the voltage difference between membrane depolarization baseline and the hyperpolarization peak after the first AP. Refer to Figure 2C for a schematic representation of these calculations. To evaluate changes in neuronal excitability related to K_v_1.1 channel kinetics, the delay of the first AP and the slope of the depolarizing ramp at the subthreshold step were analyzed: first AP delay was calculated as time from the start of current injection to the onset of the first AP (Figure 6A, upper trace). Depolarizing ramp slope was measured over a 400 ms time window from the end of the current injection at the subthreshold step before the rheobase (Figure 6A, bottom trace).

#### mEPSCs analysis

The final five minutes of continuous mEPSCs recordings were analyzed. All traces were low-pass filtered at 2 kHz and baselines adjusted. mEPSCs were detected using a template detection method with a minimum amplitude threshold of 7 pA, followed by visual verification.

#### Spontaneous network activity

LFP recordings of network activity were used. The last five minutes of recordings were analyzed for amplitude and inter-event intervals between spikes using the threshold method in Clampfit software after manual adjustment of the baseline.

### Morphological reconstruction

Following mEPSCs recordings, slices with biocytin-filled (0.2%) pyramidal neurons were fixed in 4% paraformaldehyde (PFA) for 24 h at 4°C to preserve cellular morphology. Fixed slices were washed in phosphate-buffered saline (PBS), incubated with Streptavidin Alexa-Fluor 488 conjugate secondary antibody for 2 h, washed again in PBS and mounted on glass slides covered with mounting medium (Ibidi GmbH, Gräfelfing, Germany; 50001). A coverslip was used from top to prevent drying. Labeled neurons were imaged using a confocal microscope (LSM 700, Carl Zeiss, Jena, Germany), images were acquired with Zen software (RRID: SCR_013672). Z-stacks were digitally reconstructed using the SNT plugin (RRID:SCR_016566) of the ImageJ-Fiji software (RRID: SCR_002285) for semiautomated tracing of neuronal dendritic branches^52^.

### Materials

The selective K_v_1.1 channel blocker dendrotoxin-K (DTX-K, D4813) and biocytin (B4261) were purchased from Sigma-Aldrich, Munich, Germany. The synaptic blockers NBQX (#1044) and APV (#0106/1), and the sodium channels blocker tetrodotoxin (TTX, #1078) were obtained from Tocris, Bristol, UK. SR 95531 hydrobromide (Gabazine, HB0901) was purchased from Hello Bio, Bristol, UK. Streptavidin Alexa-Fluor 488 (S11223) was used as secondary antibody to label the biocytin-filled neurons and was purchased from Thermofisher Scientific, Waltham, MA, United States.

### Statistical analysis

Statistical analysis was conducted with GraphPad Prism 5 software (GraphPad Software Inc., San Diego, CA, USA, RRID:SCR_002798) for group sizes of n ≥ 5 referring to number of cells analyzed as biological samples. Variability between human tissue slices caused occasional differences in group sizes. Normality was assessed using D’Agostino and Pearson omnibus normality test or Kolmogorov–Smirnov normality test (when n < 8). Normally distributed data were analyzed using one-way analysis of variance (ANOVA) with *post hoc* Tukey’s test for multiple comparisons. However, *post hoc* tests were performed only when ANOVA F-value was significant and Bartlett’s test showed homogeneity of variances. Comparisons between passive and active properties of pyramidal neurons at 0-3h post slicing and after 18-24 h incubation were carried out using unpaired t-tests with Welch’s correction. Statistical significance was set at p ≤ 0.05. If not otherwise specified, normally distributed data are shown as mean ± standard deviation (SD).

## Results

### Intrinsic properties of acute and cultured human CA3 pyramidal neurons

To assess the functional integrity of human hippocampal CA3 pyramidal neurons in culture, we evaluated their biophysical properties using whole-cell patch clamp recordings in current clamp configuration. Single-cell recordings were performed on 30 neurons from 11 patients, with 6 recorded in acute conditions (0-3 h after recovery) and 24 recorded after 18-24 h of incubation in culture. Both passive and active properties of these neurons remained consistent, reflecting long-term stability under experimental conditions. The resting membrane potential, measured in the first minute after establishing whole-cell configuration, was -60.4 ± 5.4 mV after 18-24 h in culture (Figure 2B).

The intrinsic firing frequency of CA3 neurons was 10 ± 1.5 Hz (mean ± SEM) at 600 pA current injection, showing a positive correlation with the amount of injected current (r^2^ = 0.9723, Figure 2A), aligning with results from rodent studies on ventral CA3 pyramidal neurons^53^. The latency of the first action potential at rheobase was 235.7 ± 146.9 ms (Figure 2A) with considerable variability, possibly due to differential K_v_1.1 channel expression across CA3 pyramidal neurons^54,55^. We further analyzed passive properties, including input resistance, rheobase, membrane time constant, voltage sag, and sag ratio (Figure 2B and Table 1). Notably, the voltage sag as well as the sag ratio showed higher variability, averaging -3.7 ± 1.9 mV and 0.18 ± 0.08 respectively. Active AP properties included amplitude and threshold, half-width, rise and decay times, and fast afterhyperpolarization (fAHP). Human CA3 neurons displayed robust action potentials with a mean threshold of -42.9 ± 5.6mV, an amplitude of 101.5 ± 6.5 mV, and a half-width of 1.23 ± 0.2 ms (Figure 2C and Table 2). These properties exhibited low variability, indicating that the neurons maintained their excitability levels over the incubation period. A comparison of the intrinsic biophysical properties between acutely (0-3 h) recorded neurons and those recorded after 18-24 h in culture revealed that acutely recorded CA3 cells were slightly more excitable with regard to input-output properties, resting membrane potential and rheobase. These results suggest an effect of recent manipulation during slice preparation (Supplementary figure 1). Overall, our findings demonstrate that fundamental passive and active biophysical properties of human CA3 pyramidal neurons remain stable after 18-24 h of incubation in culture conditions, underscoring their functional integrity and suitability for further experimental manipulations.

**Table 2:** analyzed active properties of human CA3 pyramidal neurons after 0-3 h from recovery and 18-24 h incubation in untreated control conditions. Data are expressed as mean ± SD.

| Active properties |  | 0-3 h | 18-24 h |
| --- | --- | --- | --- |
| AP amplitude (mV) | Mean | 106.3 | 101.5 |
|  | SD | 6.1 | 6.5 |
| AP threshold (mV) | Mean | -49.9 | -42.9 |
|  | SD | 3.5 | 5.6 |
| AP half width (mV) | Mean | 1.20 | 1.23 |
|  | SD | 0.2 | 0.2 |
| AP rise time (ms) | Mean | 0.28 | 0.30 |
|  | SD | 0.03 | 0.06 |
| AP decay time (ms) | Mean | 1.75 | 1.79 |
|  | SD | 0.3 | 0.3 |
| fAHP (mV) | Mean | -12.4 | -13.9 |
|  | SD | 5.4 | 5.7 |

### Human CA3 pyramidal neurons display large AMPA currents and long dendritic projections

Next, we investigated miniature excitatory AMPA currents (mEPSCs) in CA3 pyramidal neurons (n = 6 from 3 patients) cultured for 18-24 h. Additionally, recorded cells were filled with 0.2% biocytin for subsequent reconstruction and assessment of dendritic projections (Figure 3A-B). Out of six reconstructed neurons, one was excluded from morphological analysis due to insufficient biocytin filling. The remaining five neurons exhibited intact, yet variable dendritic branches (Figure 3A-B). CA3 pyramidal neuron soma diameter averaged 18.2 ± 3 μm and the total dendritic length was 5.8 ± 3.2 mm. The total number of dendrites was 23 ± 12. Apical dendritic length averaged 4.4 ± 2.7 mm, whereas basal dendrite length averaged 1.4 ± 0.8 mm. Pharmacologically isolated mEPSCs were recorded at a holding potential of -70 mV for at least 20 min (Figure 3C-F). Human CA3 pyramidal neurons exhibited mEPSCs with large amplitudes (41.4 ± 13.9 pA) and high frequencies (inter-event interval (IEI)s of 0.91 ± 0.51 ms) (Figure 3C-F). Distribution analysis showed a wide range of mEPSCs amplitudes from 10 to 480 pA (10 pA bins). The majority (95%) of events clustered within 10 to 100 pA (Figure 3E, left panel). Inter-event intervals ranged from 0.0 to 6.45 s (0.15 s bins), with a considerable portion of events evenly distributed between 0.15 and 2.10 s (Figure 3E, right panel).

**Figure 3:**
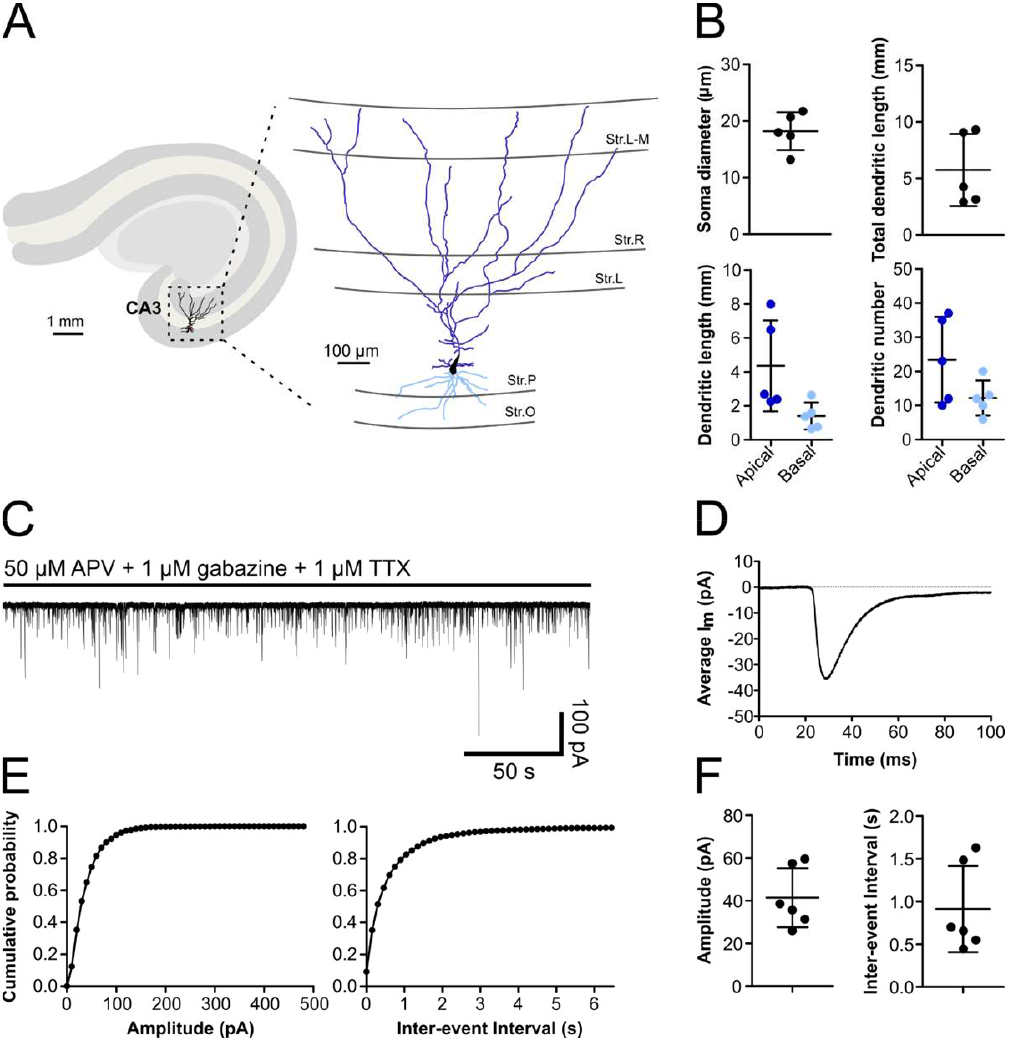
Human CA3 pyramidal neurons exhibit long dendritic branches and large spontaneous excitatory currents. **A)** left: Schematic representation of a typical hippocampal slice featuring a digitally reconstructed neuron within the CA3 subregion. Right: the same neuron at higher magnification and a view of the dendrites and their distribution across the laminar structure of the CA3 region. The apical dendritic branches are shown in dark blue, while the basal dendrites are highlighted in light blue. Abbreviations: stratum oriens (Str. O), stratum pyramidale (Str. P), stratum lucidum (Str. L), stratum radiale (Str. R), stratum lacunosum-moleculare (Str. L-M. **B)** Scatter plots with morphological properties from reconstructed pyramidal neurons (n = 5 from 3 patients). Each point represents a cell. **C)** Example trace of AMPA miniature excitatory postsynaptic currents (mEPSCs). **D)** Averaged mEPSC measured from all events during a 5 min continuous recording. **E)** Cumulative probability plots showing mEPSCs amplitude (left plot) and inter-event interval (right plot) distributions. **F)** Scatter plots depicting the mean amplitude (left) and inter-event intervals (right) of 6 pyramidal neurons from 3 patients.

### LGI1 mAb increase intrinsic excitability of human CA3 pyramidal neurons

After evaluating the biophysical properties under control conditions, we investigated the effects of LGI1 mAbs on human CA3 neurons. We compared CA3 neurons exposed to either LGI1 mAb or control antibodies for 18-24 h, alongside the effects of the K_v_1.1 channel blocker DTX-K as a pharmacological control. CA3 neurons incubated for 18-24 h with either LGI1 mAb or DTX-K exhibited a higher AP frequency compared to neurons incubated with control antibodies or left untreated (statistical analysis in Supplementary Table 2). Accordingly, both LGI1 mAb and DTX-K incubation resulted in a reduced rheobase, likewise indicating increased neuronal excitability. To explore interactions or additive effects between LGI mAb and DTX-K, a subset of slices was incubated with DTX-K and LGI1 mAb simultaneously (Figure 4, yellow): while CA3 neurons exposed to both LGI1 mAb and DTX-K showed a marked increase in AP frequency when compared to those treated with either LGI1 mAb or DTX-K alone, the rheobase was not different between LGI1 mAb or DTX-K alone and LGI1 mAb and DTX-K in combination. These results suggest that LGI1 mAb enhances the intrinsic excitability of human CA3 neurons, potentially through modulation of K_v_1.1 channel activity. The observed increase in AP frequency with combined LGI1 mAb and DTX-K treatment indicates a possible interaction between these agents, although their combined effect on rheobase remained stable.

**Figure 4:**
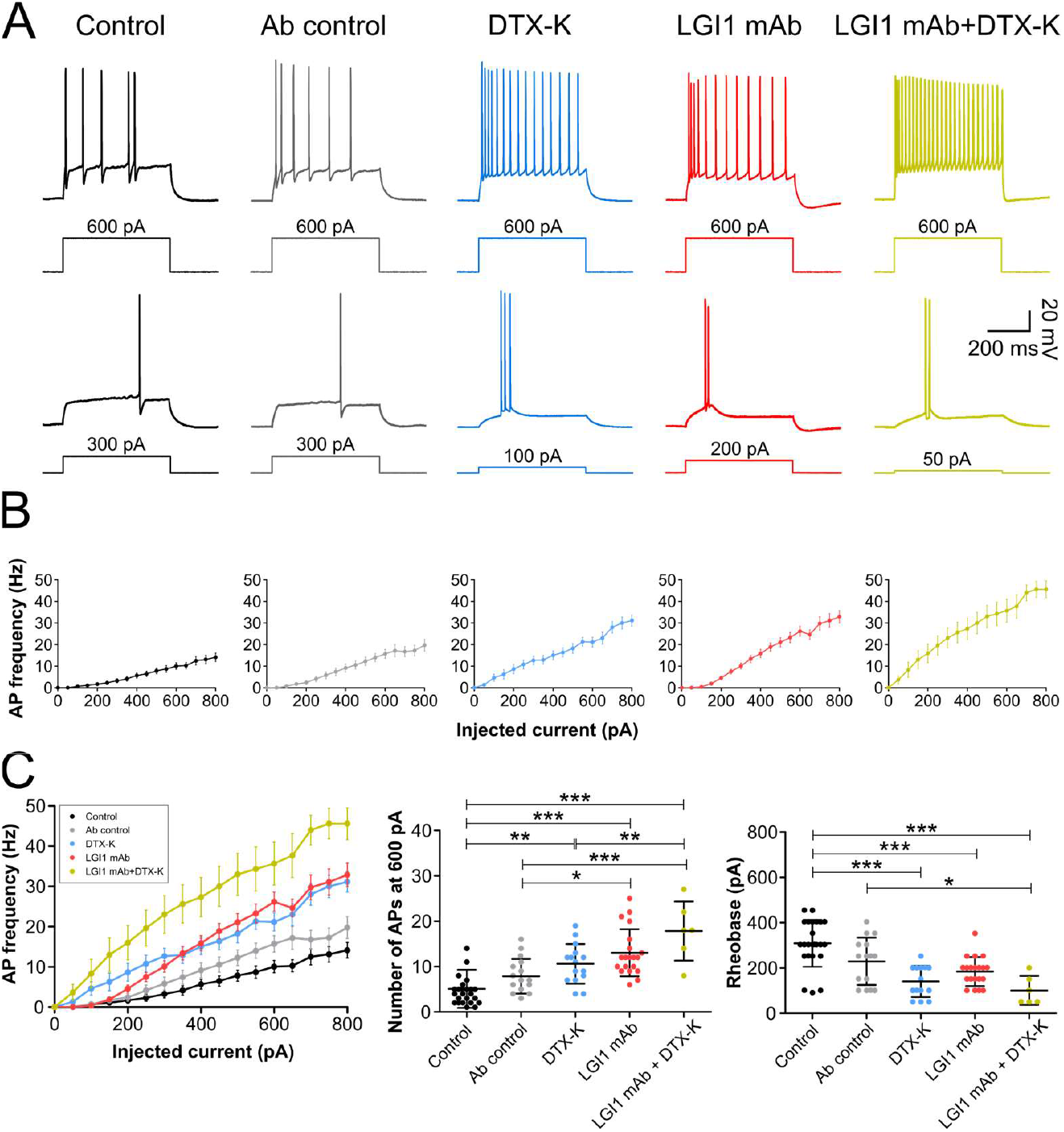
human CA3 pyramidal neurons show increased firing rate after 18-24h incubation with LGI1 mAb and DTX-K. **A)** Upper: representative AP trains evoked at 600 pA current injection in untreated controls (black), Ab control (grey), LGI1 mAb (red), DTX-K (blue), and LGI1 mAb and DTX-K combined (yellow). Lower: APs evoked at rheobase in the five conditions given above. **B)** Plots showing the I/O relation between increasing depolarizing current injections and the firing frequency. Data are shown as mean ± SEM. **C)** Left: I/O curves comparison. Middle: scatter plot with AP <u>number</u> for each condition (evoked at 600 pA; ANOVA F(4, 74) = 15.67, p < 0.0001; control: 4.7 ± 3.2, Ab control: 8.5 ± 4.5, DTX-K: 10.1 ± 4.1, LGI1 mAb: 13.1 ± 5.3, LGI1 mAb+DTX-K: 17.8 ± 6.5), right: scatter plot with <u>rheobase</u> values for each condition (ANOVA F(4, 74) = 12.91, p < 0.0001; control: 309.1 ± 103.3 pA, Ab control: 229.1 ± 104.3 pA, DTX-K: 140.7 ± 69.0 pA, LGI1 mAb: 184.4 ± 64.0 pA, LGI mAb+DTX-K: 101.1 ± 64.0 pA). Data are shown as mean ± SD. Each point represents a cell (untreated control: n = 24 from 9 patients; Ab control: n = 15 from 6 patients; DTX-K: n = 14 from 6 patients; LGI1 mAb: n = 20 from 7 patients; LGI1 mAb+DTX-K: n = 6 from 4 patients). Statistics: One-way ANOVA and Tukey’s multiple comparison test: * p<0.05; **p<0.01; ***p<0.001.

### LGI1 mAb and DTX-K alter action potential properties related to K_v_1.1 channel activity

To delve into the hypothesized reduction of K_v_1.1 channel activity induced by LGI1 autoantibodies in human CA3 pyramidal neurons, we analyzed changes in the latency of the first action potential and the slope of the depolarizing ramp at a subthreshold current step^15,54^. Both of these properties are modulated by K_v_1.1 potassium currents. For a schematic representation, refer to Figure 5A. CA3 neurons incubated with LGI1 mAb showed a significantly reduced latency of the first action potential compared to untreated controls and antibody controls (Figure 5B, left panel), whereas the reduction of latency observed with DTX-K was not significant. Additionally, the slope of the slowly depolarizing ramp at subthreshold current steps was reduced in CA3 neurons exposed to either LGI1 mAb or DTX-K, with LGI1 mAb showing a slightly greater reduction than DTX-K (Figure 5B, right panel). Concurrent exposure to LGI1 mAb and DTX-K did not induce an additional reduction in the latency of the first action potential or the slope of the depolarizing ramp compared to LGI1 mAb or DTX-K alone.

**Figure 5:**
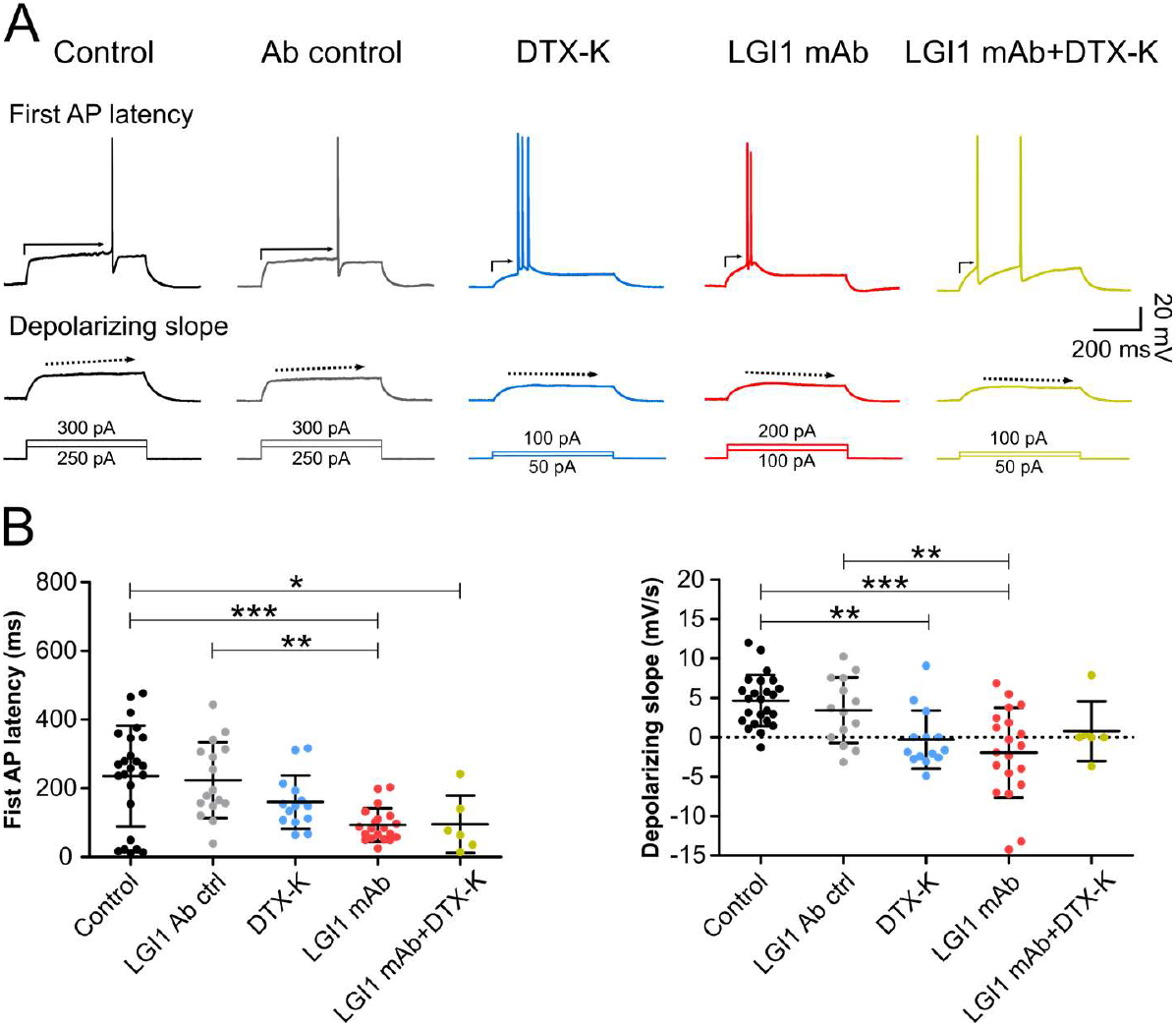
LGI1 mAb and DTX-K decreased K_v_1.1 channels activity-dependent firing properties. **A)** Upper: representative APs evoked at rheobase in five different conditions: untreated controls (black), Ab control (grey), LGI1 mAb (red), DTX-K (blue), and the combination of LGI1 mAb and DTX-K (yellow). The arrow represents the latency of the first AP. Below, representative traces showing the depolarization slope (indicated by the dashed arrow) at the subthreshold current injection. **B)** Scatter plots comparing the <u>latency of the first AP</u> (left; ANOVA F(4, 74) = 6.45, p = 0.0002; control: 235.7 ± 146.9 ms, Ab control: 223.2 ± 119.0 ms, DTX-K: 160.0 ± 77.9 ms, LGI1 mAb: 93.4 ± 48.6 ms, LGI1 mAb+DTX-K: 95.6 ± 83.5 ms) and the <u>depolarizing slope</u> (right; ANOVA F(4, 74) = 8.2, p < 0.0001; control: 4.7 ± 3.3 mV/s, Ab control: 3.6 ± 4.1 mV/s, DTX-K: -0.3 ± 3.7 mV/s, LGI1 mAb: -2.0 ± 3.7 mV/s, LGI1 mAb+DTX-K: 0.8 ± 3.8 mV/s) in the conditions given above. Data are shown as mean ± SD. Each point represents a cell (untreated control: n = 24 from 9 patients; Ab control: n = 15 from 6 patients; DTX-K: n = 14 from 6 patients; LGI1 mAb: n = 20 from 7 patients; LGI1 mAb+DTX-K: n = 6 from 4 patients). <u>Statistics</u>: One-Way ANOVA and Tukey’s multiple comparison test between groups. * p<0.05; **p<001; ***p<0.0001.

**Figure 6:**
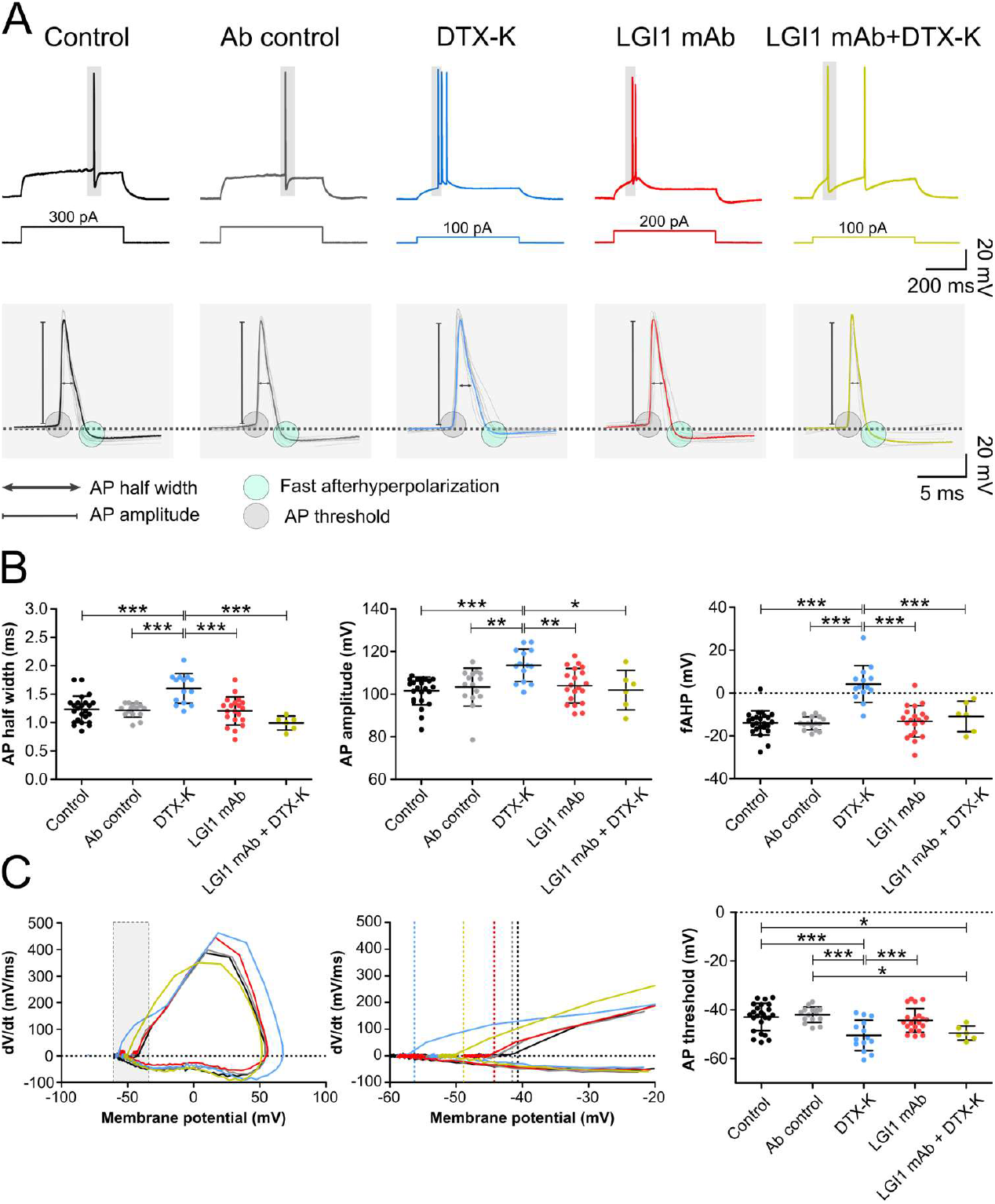
DTX-K but not LGI1 mAb affects action potential properties. **A)** Upper: representative action potentials evoked at rheobase in five different conditions: untreated control (black), Ab control (grey), LGI1 mAb (red), DTX-K (blue), and the combination of LGI1 mAb and DTX-K (yellow). Lower: highlighted APs at higher magnification showing the analyzed parameters. **B)** Scatter plots comparing AP’s <u>half width</u> (left; ANOVA F(4, 74) = 11.1, p < 0.0001; control: 1.23 ± 0.2 ms, Ab control: 1.22 ± 0.1 ms, DTX-K: 1.60 ± 0.3 ms, LGI1 mAb: 1.21 ± 0.2 ms, LGI1 mAb+DTX-K: 0.99 ± 0.1 ms), <u>amplitude</u> (middle; ANOVA F(4, 74) = 5.83, p = 0.0004; control: 101.5 ± 6.5 mV, Ab control: 103.3 ± 8.9 mV, DTX-K: 113.5 ± 7.6 mV, LGI1 mAb: 104.0 ± 8.1 mV, LGI1 mAb+DTX-K: 101.9 ± 9.3 mV), and <u>fAHP</u> (right; ANOVA F(4, 74) = 21.8, p < 0.0001; control: -13.9 ± 5.7 mV, Ab control: -14.1 ± 3.0 mV, DTX-K: 4.2 ± 8.5 mV, LGI1 mAb: -13.2 ± 7.3 mV, LGI1 mAb+DTX-K: -10.9 ± 7.1 mV). **C)** Left: AP phase plots; middle: grey-highlighted area of the phase plot is displayed at higher magnification to better appreciate the differences in threshold. Right: scatter plot showing <u>AP threshold</u> under different conditions (ANOVA F(4, 74) = 7.79, p < 0.0001; control: -42.9 ± 5.6 mV, Ab control: -42.0 ± 3.2 mV, DTX-K: -50.5 ± 6.2 mV, LGI1 mAb: -44.3 ± 4.9 mV, LGI1 mAb+DTX-K: -49.6 ± 2.9 mV). **B) and C)** Data are shown as mean ± SD. Each point represents a cell (untreated control: n = 24 from 9 patients; Ab control: n = 15 from 6 patients; DTX-K: n = 14 from 6 patients; LGI1 mAb: n = 20 from 7 patients; LGI1 mAb+DTX-K: n = 6 from 4 patients). <u>Statistics</u>: One-way ANOVA and Tukey’s multiple comparison test between groups. * p<0.05; **p<0.01; ***p<0.001.

### DTX-K and LGI1 mAb exhibit distinct effects on action potential properties

To further explore the mechanisms involved in increased neuronal excitability induced by LGI1 mAb in human CA3 neurons, we focused on the properties of the first action potential at rheobase, specifically the half width, amplitude, fast afterhyperpolarization (fAHP) and action potential threshold (Figure 6A). Our findings revealed that DTX-K altered the aforementioned action potential properties while LGI1 mAb exposure did not elicit such alterations (Figure 6B-C). Specifically, DTX-K increased the action potential half width, peak amplitude, and fAHP amplitude (Figure 6B). Additionally, DTX-K significantly decreased the action potential threshold compared to all other conditions (Figure 6C). Surprisingly, simultaneous incubation of DTX-K and LGI1 mAb did not alter action potential half width, amplitude or fAHP, suggesting that LGI1 mAb prevented DTX-K effects. However, concurrent DTX-K/LGI1 mAb exposure resulted in a significant decrease in action potential threshold, mirroring the effect observed with DTX-K alone.

### LGI1 mAb increase spontaneous network activity

Finally, since LGI1 mAb increased the excitability of individual human CA3 pyramidal neurons, we asked whether this increase goes along with increased network activity, indicative of a lower seizure threshold. To assess this, we recorded field potentials in the pyramidal layer of CA3 region (5 patients) incubated with Ab control and LGI1 mAb for 18-24 h (one slice per patient per condition). Among slices incubated with LGI1 mAb, slices from three patients exhibited increased spontaneous bursting activity following LGI1 mAb exposure (Supplementary figure 2). Due to the limited sample size, no statistical analysis was performed on these experiments.

## Discussion

In the present study, we investigated the basal passive and active biophysical properties of human CA3 pyramidal neurons, focusing specifically on the effects of human pathogenic LGI1 monoclonal autoantibodies: properties of CA3 neurons in hippocampal slices from temporal lobe patients were (a) largely consistent and retained their excitability over the incubation period, and (b) were only slightly altered after 18-24 h in culture; (c) human CA3 neurons received strong excitatory input, evidenced by large amplitudes and high frequency of spontaneous synaptic events; (d) LGI1 mAb increased the firing frequency and (e) decreased the rheobase, with an effect size similar to full K_v_1.1 channel block; (f) however, unlike K_v_1.1 channel block, LGI1 mAb only slightly altered action potential properties; finally (g) LGI1 mAb incubation was associated with increased spontaneous network activity in the CA3 network.

Our results demonstrate that human CA3 pyramidal neurons retain robust firing properties and functional integrity after 18-24 hours of culture. Passive properties of human CA3 neurons are similar to rodent CA3 pyramidal neurons^54,56–58^, but with a lower input resistance along the longitudinal axis, identified in the ventral hippocampus^53,59^. Data on voltage sag and sag ratio were highly variable, indicating inhomogeneous HCN channel expression and kinetics in human CA3 pyramidal neurons, which in part mirrors the conditions in rodent brain^60,61^. Analysis of the active properties revealed that human CA3 neurons have a slightly lower action potential threshold compared to rodents, but a higher amplitude and stronger afterhyperpolarization^53,57^. Our results align with a recent study comparing human and rodent hippocampal CA3 neurons and their connectivity^62^. Watson at al. showed that, despite similar single-cell properties, the human hippocampus is not simply a scaled version of the rodent hippocampus. Instead, it reveals sparse connectivity with high synaptic reliability, which likely enhances the auto-associative power of the hippocampus. The variability of action potential latency suggests, again, heterogeneity within human CA3 pyramidal neurons, including variations in K_v_1.1 channel expression previously documented in rodents^54,55^. Our findings align with Hemond et al., showing that passive human CA3 properties are not positively correlated with action potential latencies (data not shown), indicating the need for further investigation into CA3 pyramidal cell variability in the human hippocampus.

By comparing acute slices and short-term organotypic cultures, our study offers insights into how biophysical properties of human neurons evolve in culture and provides a foundation for future investigations in organotypic cultures. While most active and passive properties of pyramidal neurons remained consistent between acute (0-3 hours) and chronic (18-24 hours) culture conditions, neurons cultured for 18-24 hours were less excitable, exhibiting reduced action potential frequency, lower rheobase and a more negative resting membrane potential. These findings contrast with previous reports showing increased neuronal firing of cortical pyramidal neurons after 2-3 days *in vitro*^49^ and dentate gyrus granule cells incubated for 24-48 hours^63^, as well as >10 days *in vitro*^64^. However, the study from Le Duigou et al. also showed that human CA2/CA3 pyramidal neurons exhibited a marked increase in rheobase and decreased firing activity after 20-41 days *in vitro* compared to acute conditions (supplementary material in ref.^64^). The discrepancy in excitability changes over time in different brain regions may be due to differences in homeostatic and short-term plasticity induced by slicing and culture conditions^65^. These mechanisms, including neurite outgrowth, reactive synaptogenesis, and mossy fiber sprouting in the CA3 area, may contribute to the formation of new functional synaptic contacts and the restoration of physiological transmission^66–68^. Our results suggest that neuronal properties change in organotypic slice cultures, an important aspect to be considered when studying human neuronal physiology in *ex vivo* slices.

Human CA3 neurons exhibited spontaneous excitatory (AMPA) currents with notably large amplitudes and frequencies. The high average number of mEPSCs with amplitudes > 100 pA amplitudes suggests the existence of “giant” AMPA mEPSCs in human CA3 neurons, a phenomenon previously reported in rodents^69,70^. Studies in the rat have indicated that these large amplitude currents are mediated by single large vesicles at the mossy fiber to CA3 pyramidal cell synapses^69^. The conservation of mossy fibers properties and hallmarks similarities have been recently described between human and rodent hippocampus^71^. Due to the lack of proper controls for human hippocampi, however, it is difficult to determine whether these phenomena are reflecting a physiological condition or are due to altered neuronal properties in an epileptic hippocampal tissue long-term exposed to anti-seizure medication.

A critical point in our study was the observation of short total dendritic lengths of CA3 pyramidal neurons. These measurements contrast with the expected extensive dendritic branching observed in human cortical and hippocampal CA1 pyramidal neurons^36,72^. This discrepancy might be attributed to the low sample size and high variability in hippocampal CA3 pyramidal neuron morphology. Although the slice thickness allowed for the inclusion of all dendrites extending into the depth of the slice, basal and apical dendritic branch lengths may vary depending on their orientation relative to the slice surface. This aspect is particularly important for the recorded and filled pyramidal neurons in the superficial part of the slices, where alignment variations could significantly impact the measured dendritic length^45^. Further research with larger sample sizes is necessary to confirm these findings and understand the underlying causes of this variability.

Differences in neuronal properties between animal models and humans pose challenges in translating findings from animal models to clinical trials^37–42,73^. We therefore investigated the effects of human LGI1 mAb in human hippocampal slices. In line with previous studies on rodent models, exposure to LGI1 mAb increased neuronal firing frequency and intrinsic excitability of human CA3 pyramidal neurons in absence of synaptic input, mirroring the effects observed with the K_v_1.1 channel blocker DTX-K. This increase in excitability likely results from a decreased K_v_1.1 channel activity at the AIS, supported by alterations in action potential properties indicative of K_v_1.1 channel modulation: the latency of the first action potential and the ramp slope at subthreshold depolarizing currents^9,15,54^. Despite a decreased action potential latency, the lack of differences between control and DTX-K exposure could be due to variability among neurons in untreated conditions.

Our study highlights distinct effects of LGI1 mAb and DTX-K on action potential properties that further emphasize the specificity of LGI1 autoantibodies in modulating neuronal function. While LGI1 mAb increased firing activity, reduced latency of the first action potential and depolarizing ramp slope, DTX-K induced broader changes in action potential characteristics, including half width, amplitude, threshold and fast afterhyperpolarization. Concurrent exposure to LGI1 mAb and DTX-K revealed complex interactions, with LGI1 mAb preventing some effects of DTX-K while synergistically influencing action potential threshold. This suggests a saturation effect on K_v_1.1 channels, where combined action fails to elicit additional changes beyond those observed individually. The absence of LGI1 mAb effects on the main action potential properties is unexpected, as K_v_1.1 channels play a pivotal role not only in the timely generation of the action potential but also in shaping its waveform^74^. However, this observation is consistent with findings from a recent study that examined the impact of this autoantibody on rat hippocampal slice cultures^23^.

The discrepancies between LGI1 mAb and DTX-K on action potential properties likely stem from their distinct molecular targets and mechanisms of action. LGI1 mAb appears to interfere with the protein complex involving LGI1, ADAM22, and K_v_1.1 channels at the AIS, leading to K_v_1.1 channel internalization^9,22,23^. The specific interaction between Kv1.1 and LGI1 mAb or DTX-K, respectively, could lead to different responses with regard to homeostatic plasticity or intrinsic excitability, which was previously suggested by K_v_1.1 channel expression in rodent hippocampal CA3 and dentate gyrus cells^16,74,75^. The hyperexcitability resulting from K_v_1.1 channel internalization by LGI1 mAb might trigger a regulatory process involving other receptors and channels, such as a reduction in the density of voltage-gated sodium channels at the AIS, as previously shown in chronically depolarized hippocampal neurons^76^. This regulatory response could ultimately lead to a decrease in neuronal activity. In our study, this phenomenon could explain why we did not observe alterations in action potential threshold and other action potential properties after 18-24 hours of exposure to LGI1 mAb.

The observation of spontaneous bursting network activity in human hippocampal slices following LGI1 mAb exposure underscores the broader impact of autoantibodies on neuronal network function. Spontaneous ictal-like network activity without pharmacological or electrical manipulation occurs rarely^77,78^, potentially due to cut axonal connections in a neuronal network with lower cellular density than in rodents. Only interictal-like spontaneous activity can be observed more frequently^79,80^. Our finding, to our knowledge never observed in human CA3 *in vitro* preparations before, highlights the importance of studying autoimmune-mediated alterations in neuronal excitability at the network level, focusing on species-specific effects.

Our findings suggest that human brain tissue obtained from epilepsy patients can be used to study brain pathologies beyond epilepsy. One concern is that the intrinsic properties of neurons from epilepsy patients may be altered compared to neurons from a healthy brain, potentially causing an excitation-inhibition imbalance and increased network excitability^81^. However, spontaneous epileptiform activity has not been observed in single cortical or hippocampal neurons from epilepsy patients^82–84^. The difference between untreated controls, which lack spontaneous activity, and the presence of ictal-like activity after incubation with LGI1 autoantibodies, along with increase in CA3 neurons intrinsic excitability, indicates that tissue resected from patients retains physiological properties and allows to study changes upon pathological perturbations, such as LGI1 mAb exposure used in our study.

## Conclusions

The functional characterization of human CA3 pyramidal neurons in the presence of LGI1 autoantibodies in our study leverages the advantages of *ex vivo* human brain tissue and underscores the importance of human-specific investigations. Despite limitations related to sample size, *ex vivo* models and tissue variability, our findings pave the way for future investigation of human pathologies using human organotypic brain slice cultures. Our approach enables the investigation of pharmacological compounds, mechanisms of neurodegenerative diseases, and also autoantibodies, using human brain tissue obtained from drug-resistant epilepsy patients. Additionally, the observed variability in dendritic morphology and action potential properties underlines the importance of considering individual neuron heterogeneity in studies of human brain tissue. Future investigations should aim to expand sample sizes and address these variabilities, providing a deeper understanding of how disease-associated antibodies like LGI1 mAb affect human neuronal function and excitability at both the cellular and network levels.

## Acknowledgments

We are deeply grateful to the patients for donating the tissue and to the surgical teams across all involved neurosurgery departments for their availability and helpfulness during the sample collection and handling. We would like to thank Mandy Marbler-Pötter and Andrea Wilke (both Charité-Unversitätsmedizin, Berlin) for their excellent technical assistance. We also extend our thanks to all the researchers of the Human Brain Tissue group of Charité Berlin for their invaluable support.

