## Supplementary information for "The functional impact of LGI1 autoantibodies on human CA3 pyramidal neurons"

**Supplementary table 1**: patients' information.

| Sample ID | Diagnosis | Age | Biological sex | Medication 1 (mg) | Medication 2 (mg) | Medication 3 (mg) |
|---|---|---|---|---|---|---|
| #1 | Temporal lobe epilepsy | 27 | Male | Lacosamide (400) | - | - |
| #2 | Temporal lobe epilepsy | 52 | Male | Lamotrigine (300) | - | - |
| #3 | Temporal lobe epilepsy | 33 | Male | Lamotrigine (800) | Brivaracetam (200) | - |
| #4 | Temporal lobe epilepsy | 52 | Male | Levetiracetam (3000) | Primidone (750) | Carbamazepine (1200) |
| #5 | Temporal lobe epilepsy | 60 | Male | Oxcarbazepine (1200) | Levetiracetam (3000) | - |
| #6 | Temporal lobe epilepsy | 22 | Male | Oxcarbazepine (1500) | Valproic acid (1050) | Brivaracetam (225) |
| #7 | Temporal lobe epilepsy | 42 | Female | Levetiracetam (2000) | Clobazam (3) | - |
| #8 | Temporal lobe epilepsy | 51 | Female | Brivaracetam (200) | Lamotrigine (150) | Valproic acid (1500) |
| #9 | Temporal lobe epilepsy | 57 | Male | Gabapentin (900) | Lamotrigine (600) | - |
| #10 | Temporal lobe epilepsy | 26 | Male | Lacosamide (600) | Levetiracetam (1500) | - |
| #11 | Temporal lobe epilepsy | 13 | Male | Lacosamide (250) | Brivaracetam (75) | Midazolam (if necessary) |
| #12 | Temporal lobe epilepsy | 22 | Male | Lacosamide (250) | Oxcarbazepine (1575) | - |
| #13 | Temporal lobe epilepsy | 51 | Female | - | - | - |
| #14 | Temporal lobe epilepsy | 40 | Male | Lacosamide (100) | Brivaracetam (100) | |

**Supplementary Table 2**: statistical results of firing frequency in the five different conditions. Two-way ANOVA interaction F(4, 72) = 5.41, p-value < 0.0001.

| | Condition | | Bonferroni *post hoc* test A vs B | |
|---|---|---|---|---|
| **Injected current** | **A** | **B** | **t-value** | **p-value** |
| 200 pA | Control | Ab control | 0.327 | p>0.05 |
| 200 pA | Control | DTX-K | 3.117 | p>0.05 |
| 200 pA | Control | LGI1 mAb | 1.418 | p>0.05 |
| 200 pA | Control | LGI1 mAb + DTX-K | 4.590 | p<0.001 |
| 200 pA | Ab control | DTX-K | 2.446 | p>0.05 |
| 200 pA | Ab control | LGI1 mAb | 0.908 | p>0.05 |
| 200 pA | Ab control | LGI1 mAb + DTX-K | 4.052 | p<0.01 |
| 200 pA | DTX-K | LGI1 mAb | 1.735 | p>0.05 |
| 200 pA | DTX-K | LGI1 mAb + DTX-K | 2.212 | p>0.05 |
| 200 pA | LGI1 mAb | LGI1 mAb + DTX-K | 3.568 | p<0.01 |
| 300 pA | Control | Ab control | 1.125 | p>0.05 |
| 300 pA | Control | DTX-K | 4.188 | p<0.001 |
| 300 pA | Control | LGI1 mAb | 3.312 | p>0.05 |
| 300 pA | Control | LGI1 mAb + DTX-K | 6.320 | p<0.001 |
| 300 pA | Ab control | DTX-K | 2.670 | p>0.05 |
| 300 pA | Ab control | LGI1 mAb | 1.774 | p>0.05 |
| 300 pA | Ab control | LGI1 mAb + DTX-K | 5.119 | p<0.001 |
| 300 pA | DTX-K | LGI1 mAb | 1.095 | p>0.05 |
| 300 pA | DTX-K | LGI1 mAb + DTX-K | 3.117 | p>0.05 |
| 300 pA | LGI1 mAb | LGI1 mAb + DTX-K | 4.038 | p<0.01 |
| 400 pA | Control | Ab control | 1.511 | p>0.05 |
| 400 pA | Control | DTX-K | 4.188 | p<0.001 |
| 400 pA | Control | LGI1 mAb | 4.964 | p<0.001 |
| 400 pA | Control | LGI1 mAb + DTX-K | 6.940 | p<0.001 |
| 400 pA | Ab control | DTX-K | 2.323 | p>0.05 |
| 400 pA | Ab control | LGI1 mAb | 2.825 | p>0.05 |
| 400 pA | Ab control | LGI1 mAb + DTX-K | 5.432 | p<0.001 |
| 400 pA | DTX-K | LGI1 mAb | 0.356 | p>0.05 |
| 400 pA | DTX-K | LGI1 mAb + DTX-K | 3.700 | p<0.01 |
| 400 pA | LGI1 mAb | LGI1 mAb + DTX-K | 3.579 | p<0.01 |
| 500 pA | Control | Ab control | 2.459 | p>0.05 |
| 500 pA | Control | DTX-K | 4.651 | p<0.001 |
| 500 pA | Control | LGI1 mAb | 6.440 | p<0.001 |
| 500 pA | Control | LGI1 mAb + DTX-K | 8.064 | p<0.001 |
| 500 pA | Ab control | DTX-K | 2.345 | p>0.05 |
| 500 pA | Ab control | LGI1 mAb | 3.685 | p<0.01 |
| 500 pA | Ab control | LGI1 mAb + DTX-K | 6.185 | p<0.001 |
| 500 pA | DTX-K | LGI1 mAb | 1.209 | p>0.05 |
| 500 pA | DTX-K | LGI1 mAb + DTX-K | 4.444 | p<0.001 |
| 500 pA | LGI1 mAb | LGI1 mAb + DTX-K | 3.725 | p<0.01 |
| 600 pA | Control | Ab control | 2.459 | p>0.05 |
| 600 pA | Control | DTX-k | 4.961 | p<0.001 |
| 600 pA | Control | LGI1 mAb | 7.829 | p<0.001 |
| 600 pA | Control | LGI1 mAb + DTX-K | 8.200 | p<0.001 |
| 600 pA | Ab control | DTX-K | 2.151 | p>0.05 |
| 600 pA | Ab control | LGI1 mAb | 4.384 | p<0.001 |
| 600 pA | Ab control | LGI1 mAb + DTX-K | 5.958 | p<0.001 |
| 600 pA | DTX-K | LGI1 mAb | 2.133 | p>0.05 |
| 600 pA | DTX-K | LGI1 mAb + DTX-K | 4.364 | p<0.001 |
| 600 pA | LGI1 mAb | LGI1 mAb + DTX-K | 2.963 | p>0.05 |

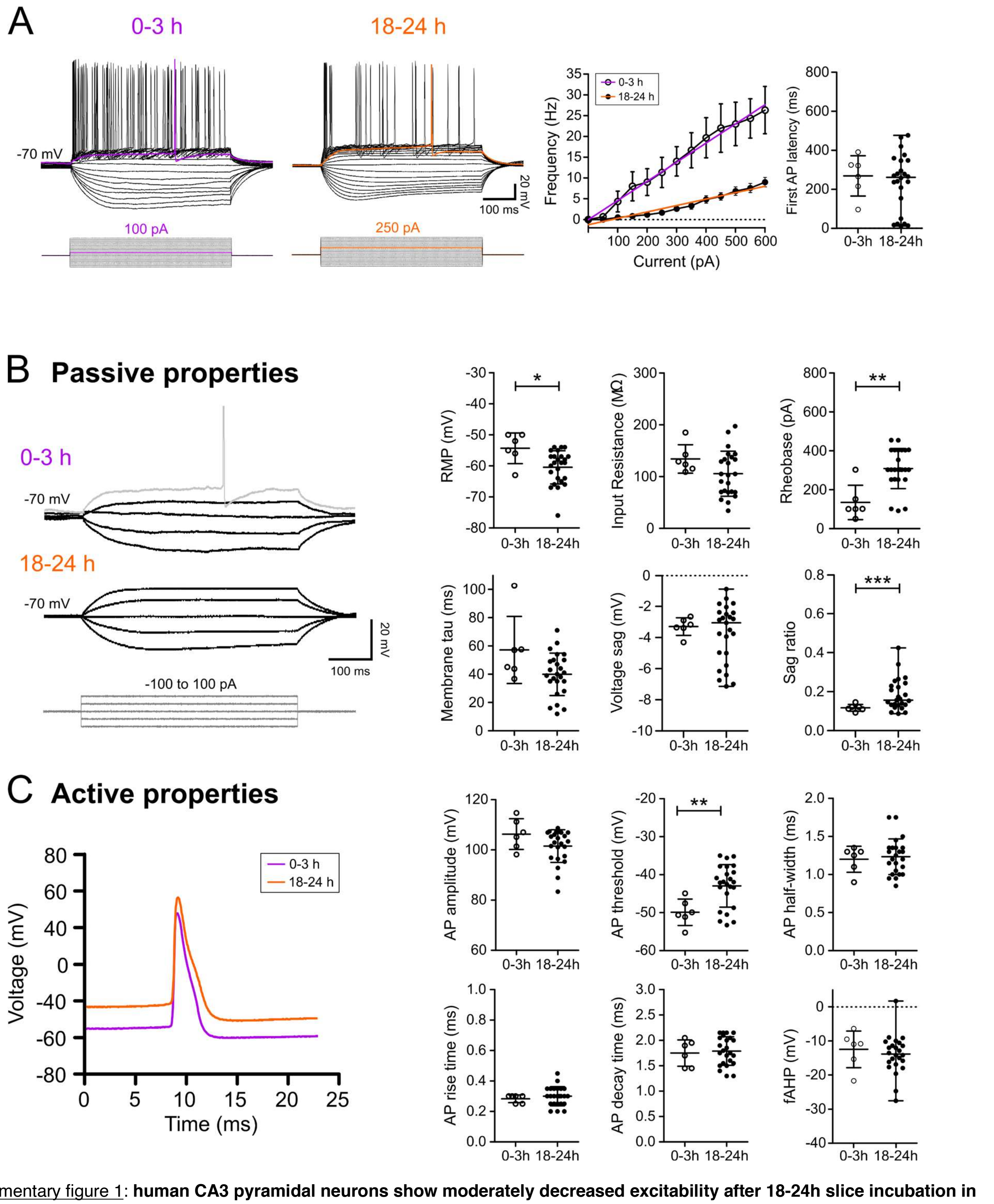


Supplementary figure 1: **human CA3 pyramidal neurons show moderately decreased excitability after 18-24h slice incubation in control conditions compared to acute slices**. **A)** Left: representative traces of trains of action potentials, induced by increasing hyperpolarizing and depolarizing current injections (from -400 to 600 pA, 50 pA steps) obtained after 0-3h from recovery and 18-24h incubation, respectively. The first action potential (AP) evoked at the rheobase is highlighted in purple (0-3h) or red (18-24 h). In the middle, the plot shows the input-output linear relation between increasing depolarizing current injections and the neuronal firing frequency after 0-3h from recovery (empty points and purple line) and 18-24h incubation (black points and orange line) in control conditions (Data shown as mean ± SEM). On the right, a scatter plot showing the first AP latency after 0-3h and 18-24h. Data are shown as mean ± SD, 0-3h n = 6 from 2 patients; 18-24h n = 24 from 9 patients. Cells recorded acutely were more excitable than after 18-24 h (0-3h: r2 = 0.9896, 18-24h: r2 = 0.9494; ANCOVA F(1, 22) = 294.6, p-value < 0.0001). **B)** Left: the experimental steps used to determine the passive properties. Right: scatter plots comparing the main passive properties of neurons recorded after 0-3h (empty points) and 18-24h (black points). **C)** Left: representative action potentials recorded at the rheobase, which was employed to quantify the active properties. Right: scatter plots comparing the active properties of neurons recorded after 0-3h (empty points) and 18-24h (black points). **B)-C)** Each point indicates an individual cell and data are shown as mean ± SD (0-3h n = 6 from 2 patients, 18-24h n = 24 from 9 patients). Statistical comparison between the two groups was performed with unpaired t-test with Welch's correction. * p<0.05; **p<0.01; ***p<0.001.

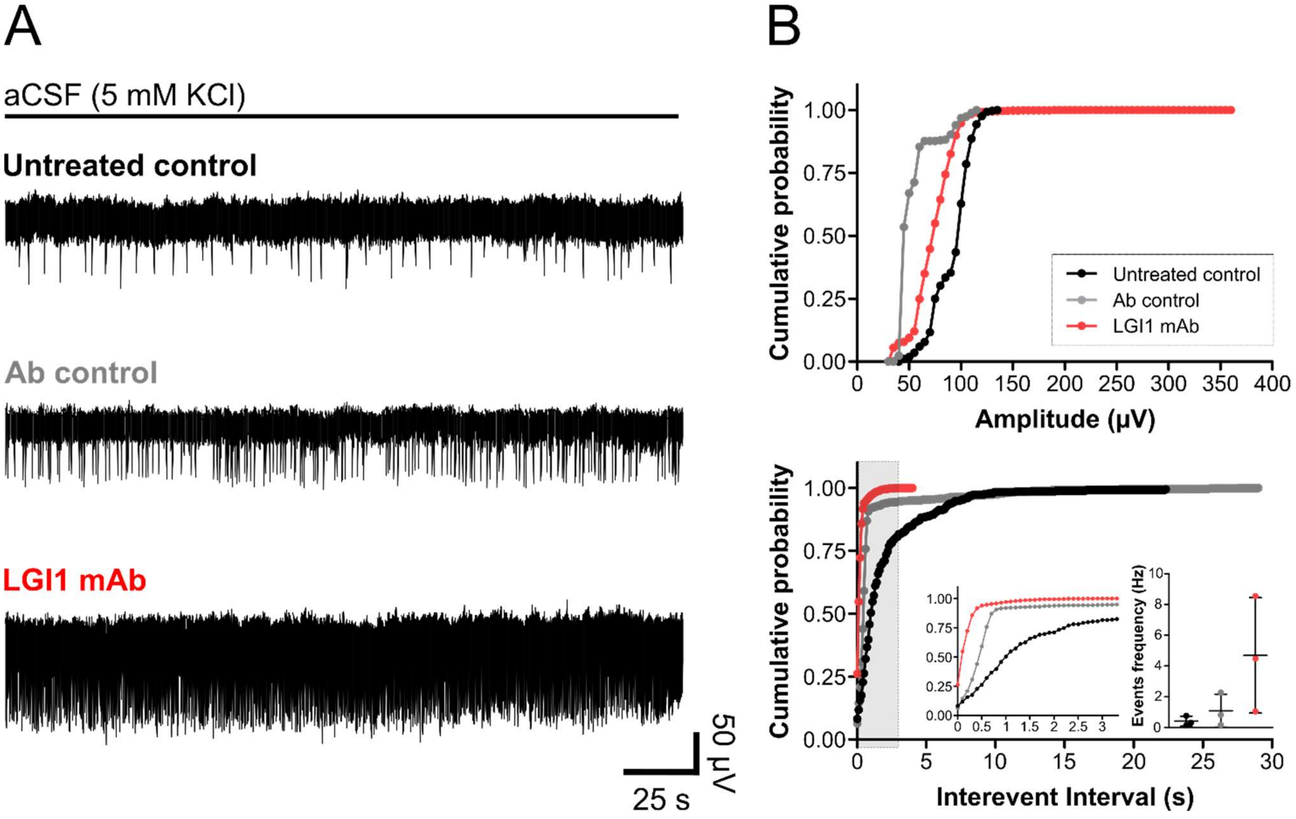


Supplementary figure 2: **LGI1 mAb increased spontaneous network activity after 24 hours of incubation**. **A)** Example traces of spontaneous network activity recorded in aCSF 5 mM KCl after 24 h of incubation in untreated controls (upper), Ab control (middle), and LGI1 mAb (bottom). **B)** Cumulative distribution plots showing spike amplitude (upper) and inter-spike interval (lower), inset: on the left, grey-highlighted area of the cumulative plot is displayed at higher magnification; on the right, scatter plot depicting the spike frequency. Untreated control condition (black), Ab control (grey) and LGI1 mAb (red). Each dot of the cumulative distributions represents a bin, while the dots in the scatter plot of spike frequency represent 5 min recording from each patient's slices incubated in the three different color-coded conditions.
